# Aging and neurodegeneration are associated with increased mutations in single human neurons

**DOI:** 10.1101/221960

**Authors:** Michael A. Lodato, Rachel E. Rodin, Craig L. Bohrson, Michael E. Coulter, Alison R. Barton, Minseok Kwon, Maxwell A. Sherman, Carl M. Vitzthum, Lovelace J. Luquette, Chandri Yandava, Pengwei Yang, Thomas W. Chittenden, Nicole E. Hatem, Steven C. Ryu, Mollie B. Woodworth, Peter J. Park, Christopher A. Walsh

## Abstract

It has long been hypothesized that aging and neurodegeneration are associated with somatic mutation in neurons; however, methodological hurdles have prevented testing this hypothesis directly. We used single-cell whole-genome sequencing to perform genome-wide somatic single-nucleotide variant (sSNV) identification on DNA from 161 single neurons from the prefrontal cortex and hippocampus of fifteen normal individuals (aged 4 months to 82 years) as well as nine individuals affected by early-onset neurodegeneration due to genetic disorders of DNA repair (Cockayne syndrome and Xeroderma pigmentosum). sSNVs increased approximately linearly with age in both areas (with a higher rate in hippocampus) and were more abundant in neurodegenerative disease. The accumulation of somatic mutations with age—which we term genosenium—shows age-related, region-related, and disease-related molecular signatures, and may be important in other human age-associated conditions.

**One-Sentence Summary:** Somatic single-nucleotide variants accumulate in human neurons in aging with regional specificity and in progeroid diseases.

## Main Text

Aging in humans brings increased incidence of nearly all diseases, including neurodegenerative diseases (1). Markers of DNA damage increase in the brain with age (2), and genetic progeroid diseases such as Cockayne syndrome (CS) and Xeroderma pigmentosum (XP), both caused by defects in DNA damage repair (DDR), are associated with neurodegeneration and premature aging (3). Mouse models of aging, CS, and XP have shown inconsistent relationships between these conditions and the accumulation of permanent somatic mutations in brain and non-brain tissue (4–7). While analysis of human bulk brain DNA, comprised of multiple proliferative and non-proliferative cell types, revealed an accumulation of mutations during aging in the human brain (8), it is not known whether permanent somatic mutations accumulate with age in mature neurons of the human brain. Here, we quantitatively examined whether aging or disorders of defective DDR results in more somatic mutations in single postmitotic human neurons.

Somatic mutations that occur in postmitotic neurons are unique to each cell, and thus can only be comprehensively assayed by comparing the genomes of single cells (9). Therefore, we analyzed human neurons by single-cell whole-genome sequencing (WGS). Since alterations of the prefrontal cortex (PFC) have been linked to age-related cognitive decline and neurodegenerative disease (10), we analyzed 93 neurons from PFC of 15 neurologically normal individuals (Tables 1, S1, S2) from ages 4 months to 82 years. We further examined 26 neurons from the hippocampal dentate gyrus (DG) of 6 of these individuals because the DG is a focal point for other age-related degenerative conditions such as Alzheimer’s disease. Moreover, the DG is one of the few parts of the brain that appears to undergo neurogenesis after birth (11), which might create regional differences in number and type of somatic mutations. Finally, to test whether defective DDR in early-onset neurodegenerative diseases is associated with increased somatic mutations, we analyzed 42 PFC neurons from 9 individuals diagnosed with CS or XP (Tables 1, S3).

**Table 1.** Case information and number of neurons analyzed in this study.

| Case ID | Sex | Age (yrs) | Diagnosis | PFC Neurons | DG Neurons |
| --- | --- | --- | --- | --- | --- |
| **Infant** |  |  |  |  |  |
| 1278 | M | 0.4 | Normal | 9 | - |
| 5817 | M | 0.6 | Normal | 4 | - |
|  |  |  |  | **13** | **-** |
| **Adolescent** |  |  |  |  |  |
| 4638 | F | 15.1 | Normal | 10 | - |
| 1465 | M | 17.5 | Normal | 22 | - |
| 5532 | M | 18.4 | Normal | 4 | 5 |
| 5559 | F | 19.8 | Normal | 5 | 5 |
|  |  |  |  | **41** | **10** |
| **Adult** |  |  |  |  |  |
| 4643 | F | 42.2 | Normal | 10 | - |
| 5087 | M | 44.9 | Normal | 4 | 5 |
| 936 | F | 49.2 | Normal | 3 | - |
|  |  |  |  | **17** | **5** |
| **Aged** |  |  |  |  |  |
| 5840 | M | 75.3 | Normal | 3 | 3 |
| 5219 | F | 77 | Normal | 4 | - |
| 5171 | M | 79.2 | Normal | 4 | - |
| 5511 | F | 80.2 | Normal | 3 | - |
| 5657 | M | 82.2 | Normal | 5 | 5 |
| 5823 | F | 82.7 | Normal | 3 | 3 |
|  |  |  |  | **22** | **11** |
| **Cockayne syndrome** |  |  |  |  |  |
| 1762 | F | 4.4 | CS (CSB) | 6 | - |
| 1124 | F | 4.7 | CS (CSB) | 3 | - |
| 1286 | M | 5.8 | CS (CSB) | 4 | - |
| 580 | F | 8.4 | CS (CSB) | 4 | - |
| 5105 | M | 8.7 | CS (CSB) | 6 | - |
| 682 | M | 32.8 | CS (CSB) | 4 | - |
|  |  |  |  | **27** | **-** |
| **Xeroderma pigmentosum** |  |  |  |  |  |
| 5379 | F | 24 | XP (XPA) | 6 | - |
| 5316 | F | 44.5 | XP (XPA) | 3 | - |
| 5416 | F | 46 | XP (XPD) | 6 | - |
|  |  |  |  | **15** | **-** |
| **Total** | 24 cases |  |  | **135 PFC Neurons** | **26 DG Neurons** |
|  |  |  |  | **161 neurons** |  |

We isolated single neuronal nuclei using flow cytometry, lysed the nuclei on ice in alkaline conditions as we previously performed (12, 13) (Methods) to minimize lysis-induced artifacts, amplified their genomes using multiple displacement amplification (MDA), and subjected the amplified DNA to 45X WGS (Figure 1A). To identify somatic single nucleotide variants (sSNVs) with high confidence, we developed a bioinformatic pipeline called Linked-Read Analysis (LiRA) (14) to delineate true double-stranded sSNVs from single-stranded variants and artifacts. This method employs read-based linkage of candidate sSNVs with nearby germline SNPs and performs a model-based extrapolation of the genome-wide mutational frequency based on the ~20% of sSNVs that are sufficiently close to germline SNPs (Figure 1B; Methods). Importantly, sSNVs determined by our algorithm (Tables S4, S5) showed alternate allele frequency distribution strikingly matching that of the germline SNVs (Figure S1). sSNV counts were not systematically influenced by technical metrics, such as post-mortem interval, time in storage, and coverage uniformity (Figure S2).

**Figure 1.**
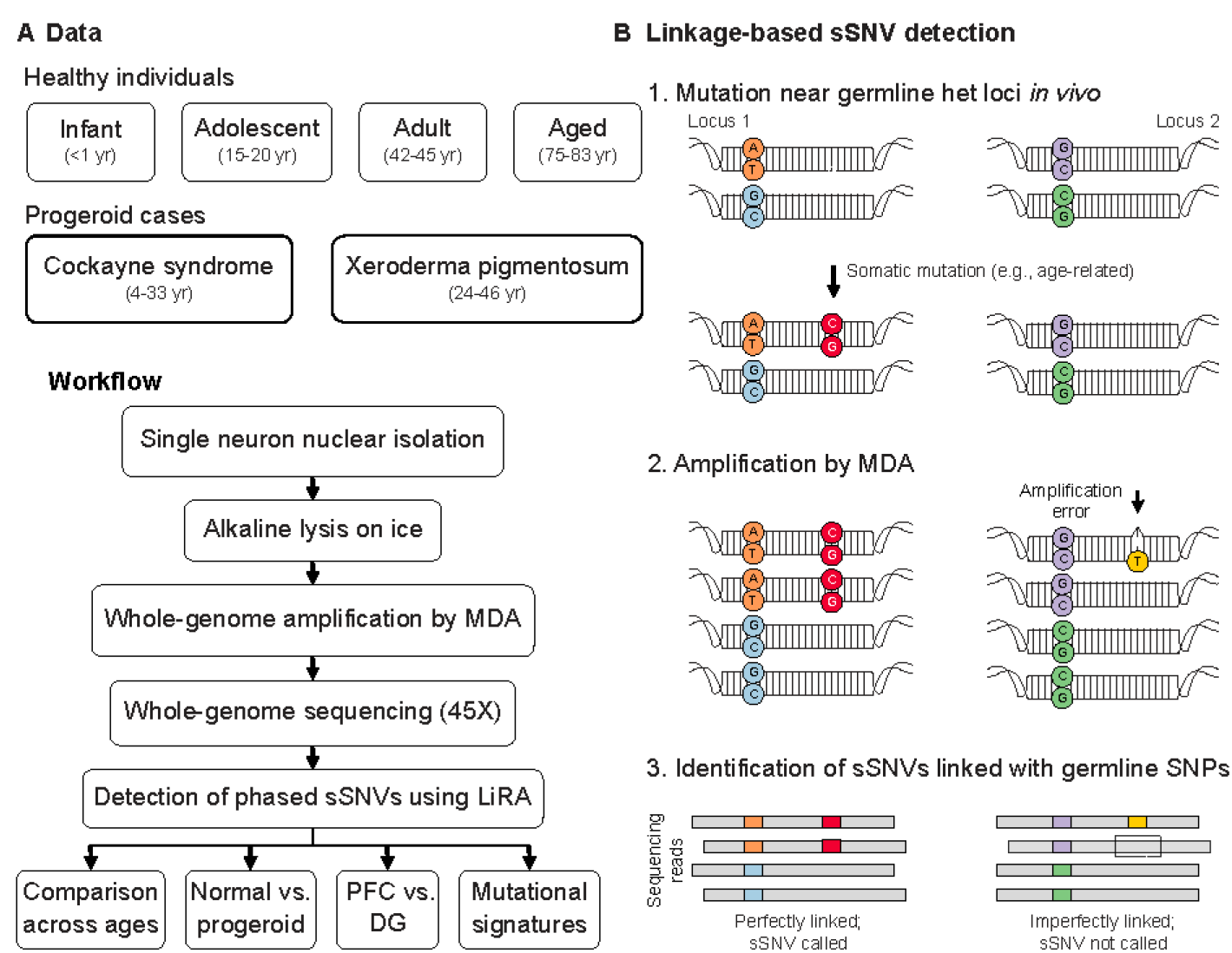
Detection of sSNVs across individuals and brain regions using single-neuron WGS and linkage-based analysis. (A) Experimental outline. (B) Schematic of linkage-based mutation calling. A somatic mutation (red) may occur on one allele (Allele 1) of a locus (Locus 1), potentially in close proximity to a germline heterozygous site (blue and light blue), while other loci, such as Locus 2, remain unmutated. Later amplification errors could create a mismatch (purple) on one strand of one allele of Locus 2 near a germline variant (light green). For Locus 1, any WGS read that covers both sites and contains the germline (blue) variant should also contain the somatic (red) variant, thus these variants are perfectly linked. In contrast, at Locus 2 some but not all reads that cover the relevant germline variant (light green) will contain the somatic “variant” candidate (purple), generating two classes of reads, some with the somatic variant on that allele, some without. Only perfectly linked sSNV candidates were considered in this study.

Across all normal neurons, genome-wide sSNV counts correlated with age (Figures 2A, S3) (p = 9×10^−12^, mixed effects model, see Methods), despite some within-individual and within-age group heterogeneity. To explore potential variation in different brain regions, matched DG and PFC neurons were sequenced in six cases (Figures 2A, S3). Our analysis uncovered region-specific sSNV accumulation with age in both PFC (p = 4×10^−5^) and DG single neurons (p = 2×10^−7^), suggesting an almost two-fold increase in the rate of accumulation in DG (~40 sSNVs/year) relative to PFC (~23 sSNVs/year) (p = 8×10^−4^). Among the six cases, two had significant increases in DG, three had nominally increased counts in DG compared to PFC, and one had a nominally higher count in PFC (Figure 2B).

**Figure 2.**
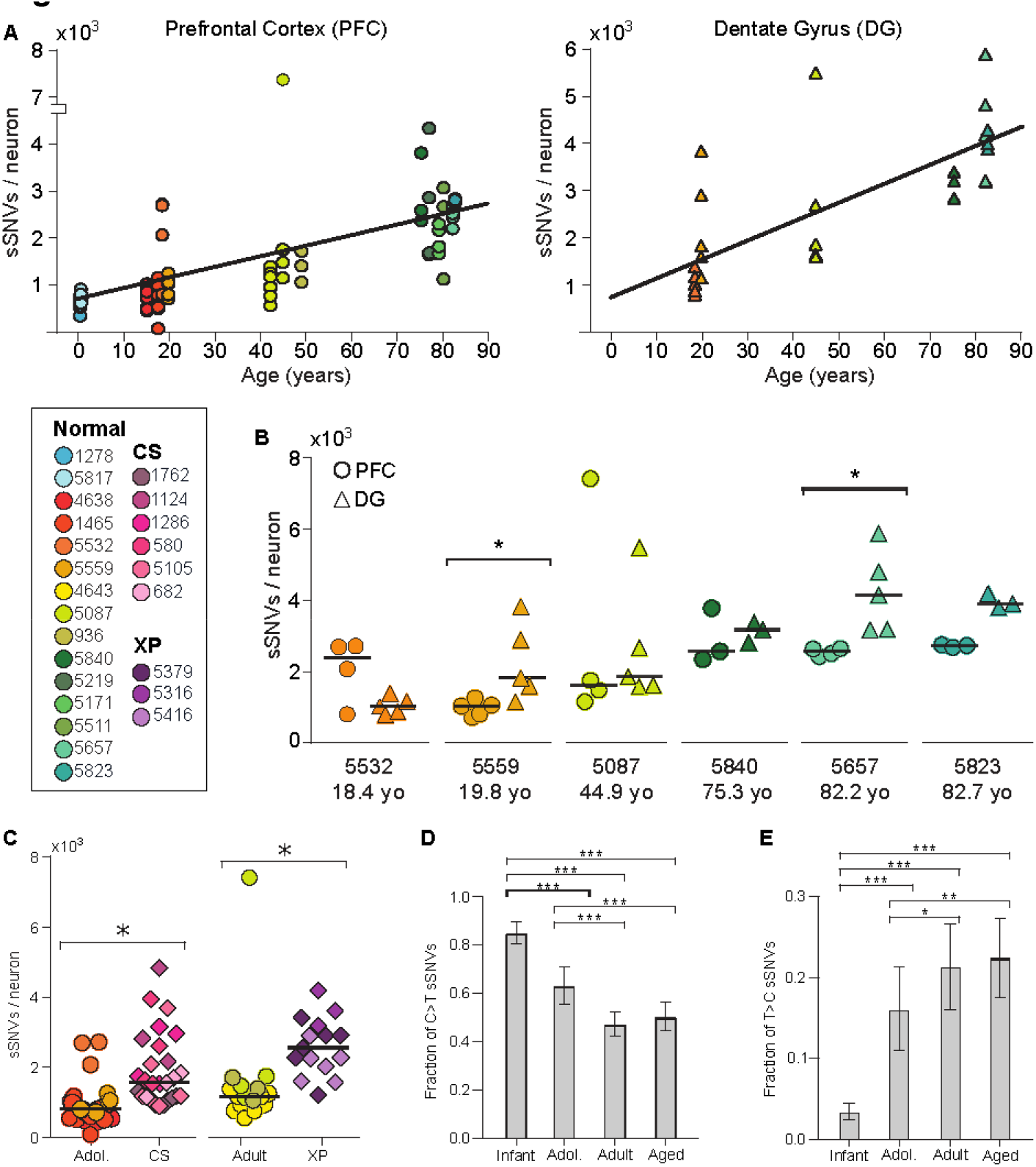
sSNVs increase with age in the PFC and DG, and are elevated in CS and XP. (A) sSNV counts plotted against age for neurons derived from PFC (circles) and DG (triangles), with linear regression lines. There is a strong correlation with age in both cases, with the rate of accumulation being nearly two-fold higher in the DG than in the PFC. (B) Comparison of sSNV counts in matched PFC and DG within brains. (C) CS and XP patient neurons display elevated sSNV counts. (D, E) Fraction of sSNVs comprised of C>T (D) and T>C (E). *, **, and *** denote p ≤ 0.05, 0.001, and 0.0001, respectively, using 2-way ANOVA with Sidak’s correction.

Neurons from postmortem brains of CS individuals showed a ~2.3-fold excess of sSNVs relative to the expected age-adjusted normal PFC rate, and XP neurons showed a ~2.5-fold increase (p = 0.006 for both) (Figure 2C). Progeroid neurons showed a similar number of sSNVs as neurons from aged normal PFC, suggesting that defective nucleotide excision repair accelerate aging via sSNV accumulation.

Molecular patterns of sSNVs also evolved with age. We previously reported that cytosine deamination influences patterns of human neuron sSNVs, resulting in abundant C>T mutations (13). C>T sSNVs accounted for most variants in the youngest PFC samples, but this fraction decreased with age (Figures 2D, S4, S5). C>T mutations, while common in many biological contexts (15–18), are also a known artifact of MDA (19). Systematic differences in C>T burden during aging suggest C>T variants are largely biological and not technical in nature. T>C variants increased in the PFC with age (Figures 2E, S4, S5), possibly representing DNA damage linked to fatty-acid oxidation (20). As demonstrated previously (13), neuronal sSNVs in normal PFC were enriched in coding exons (Figure S6, Table S6), displayed a transcriptional strand bias (Figure S7), and genes involved in neural function were enriched for neuronal sSNVs (Figure S8, Tables S7). Coincident probability modeling suggested the linear accumulation of sSNVs in our dataset would result in an exponential accumulation of biallelic deleterious coding mutations, in agreement with classical hypotheses regarding the relationship between mutation and aging (Figure S9) (21), exacerbating differences in sSNV load in aging, across brain regions, and in disease.

Mutational signature analysis (22) revealed three signatures driving single neuron mutational spectra (Figures 3A, B, S10, S11). Signature A was comprised mainly of C>T and T>C mutations and was the only signature to increase with age (p = 9×10^−12^) independent of brain region or disease status (Figure 3C-E). Signature A resembled a “clock-like” signature found in nearly all samples in a large-scale cancer genome analysis (‘Signature 5’) (22) (Figure S10). Our data show for the first time that a similar clock-like signature is also active in postmitotic cells and hence independent of DNA replication.

**Figure 3.**
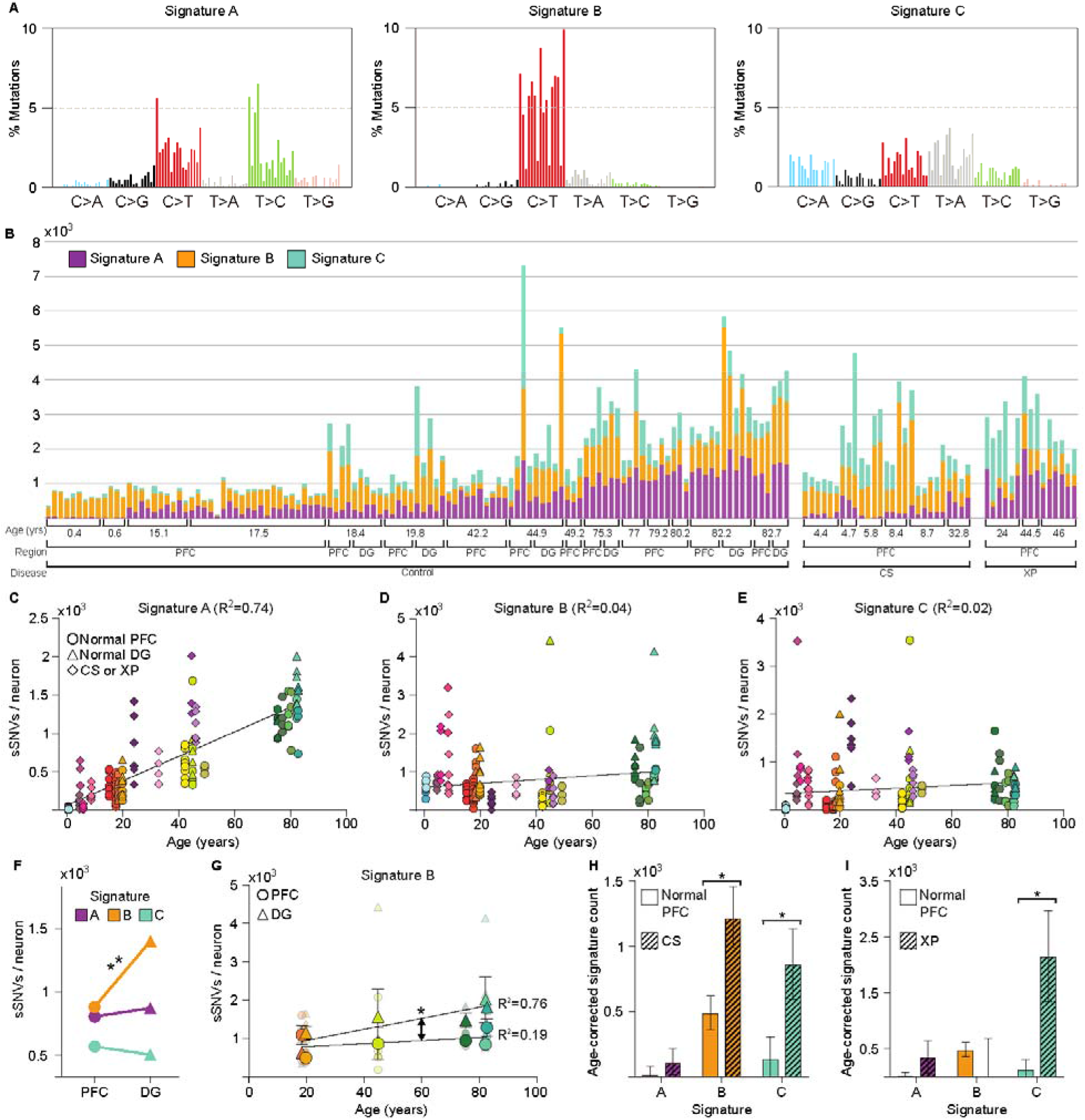
Signature analysis reveals mutational processes during aging, across brain regions, and in disease. (A) Three mutational signatures identified by matrix factorization (each substitution is classified by its trinucleotide context). (B) Number of variants from Signatures A, B, and C in each of the 161 neurons in the dataset. (C-E) Signature A strongly correlates with age, regardless of disease status or brain region, while Signatures B and C do not. (F) Signature B is enriched in DG relative to PFC neurons. (G) Signature B increased with age in DG neurons, but not in matched PFC neurons, revealing a DG-specific aging signature. Solid shapes represent regional means, and transparent shapes represent individual neurons. (H-I) Comparison of age-corrected estimate of sSNVs per signature in CS and XP compared to PFC controls revealed enrichment in Signature C in both CS and XP. * and ** denote p < 0.05 and p ≤ 0.001, respectively, mixed linear model.

Signature B consisted primarily of C>T mutations and did not correlate with age (Figure 3D), suggesting a mutational mechanism active at very young ages, perhaps prenatally. Signature B may include technical artifacts, which are primarily C>T, but bona fide clonal sSNVs are also predominantly C>T (8, 12). This signature was enriched in DG compared to PFC (p = 2×10^−4^) (Figure 3F), and increased with age in DG, but not in PFC (p = 0.04, difference in slopes) (Figure 3G). The observable difference in Signature B between these brain regions, and its correlation with age in DG alone, suggest it is dominated by a biological mechanism, and these PFC-DG differences strikingly mirror differences in neurogenesis.

A third signature, Signature C, was distinct from Signatures A and B by the presence of C>A variants, the mutation class most closely associated with oxidative DNA damage (20). Indeed, CS and XP neurons, defective in DDR, were enriched for Signature C (p = 0.016 and 0.023, respectively) (Figure 3H, I), while Signature C also increased modestly with age in normal neurons (p = 0.03). An outlier 5087 PFC neuron with the highest sSNV rate in our data set had a high proportion of Signature C mutations relative to other normal neurons, highlighting that even within a normal brain some neurons may be subject to catastrophic oxidative damage.

Our analysis revealed that sSNVs accumulated slowly but inexorably with age in the normal human brain, a phenomenon we term genosenium, and more rapidly still in progeroid neurodegeneration. Within one year of birth, postmitotic neurons already have ~300-900 sSNVs. Three signatures were associated with mutational processes in human neurons: a postmitotic, clock-like signature of aging, a possibly developmental signature that varied across brain regions, and a disease- and age-specific signature of oxidation and defective DNA damage repair. The increase of oxidative mutations in aging and in disease presents a potential target for therapeutic intervention. Further, elucidating the mechanistic basis of the clock-like accumulation of mutations across brain regions and other tissues would increase our knowledge of age-related disease and cognitive decline. CS and XP cause neurodegeneration associated with higher rates of sSNVs, and it will be important to define how other, more common causes of neurodegeneration may influence genosenium as well.

## Acknowledgements

We thank R. Sean Hill, Matthew P. Anderson, Jeffery Gulcher, Zehua Chen, Isidro Cortez, the Dana Farber Cancer Institute Flow Cytometry Core, and the Research Computing group at Harvard Medical School for assistance. Human tissue was obtained from the NIH NeuroBioBank at the University of Maryland, and we thank the donors and their families for their invaluable donations for the advancement of science. This work was supported by K99 AG054749 01, F30 MH102909, 1S10RR028832-01, T32HG002295, U01MH106883, P50MH106933, R01 NS032457 U01 MH106883, the Harvard/MIT MD-PHD program, the Stuart H.Q. and Victoria Quan Fellowship in Neurobiology, the Paul G. Allen Family Foundation. CAW is an Investigator of the Howard Hughes Medical Institute. See Supplement for full Acknowledgments.

